# Ferulic acid eicosyl ester enhances cognitive flexibility and modulates arousal-related neural circuits in mice

**DOI:** 10.64898/2026.07.20.739577

**Authors:** Dana Mayer, Iman Hassan, Firat Taskaya, Afsana Chowdhury, Evelyn Kahl, Thomas Endres, Volkmar Leßmann, Bertram Gerber, Markus Fendt

## Abstract

Cognitive deficits are a major contributor to disability in numerous neuropsychiatric and neurodegenerative disorders, yet effective pharmacological treatments remain limited. Ferulic acid eicosyl ester (FAE-20), a natural constituent of the plant *Rhodiola rosea*, has previously been identified as an enhancer of simple forms of Pavlovian conditioning in flies, bees, and mice. Here, we investigated whether FAE-20 has further potential to enhance cognitive flexibility, working memory, or spatial learning in mice, and explored potential neurobiological mechanisms underlying such enhancement. Cognitive flexibility was assessed using the attentional set-shifting task (ASST). Subchronic FAE-20 treatment significantly improved ASST performance in both male and female young adult mice, indicating enhanced cognitive flexibility. In contrast, no effects were observed on spatial working memory, assessed by spontaneous alternations in the Y-maze, or on spatial learning in the Barnes maze in either young or aged mice. Notably, FAE-20 enabled spatial learning in the Barnes maze in a subgroup of aged mice that failed to learn the task under vehicle treatment. Histological analyses using c-Fos immunohistochemistry as a marker of neural activity and doublecortin expression and spine density as markers of hippocampal plasticity revealed sex-specific effects on components of the ascending arousal system. FAE-20 increased the activation of orexinergic neurons in the lateral hypothalamus of male mice, whereas it reduced the activity of cholinergic neurons in the laterodorsal tegmental nucleus of females. No effects were detected on hippocampal neurogenesis or dendritic spine density. These findings suggest that the cognitive effects of FAE-20 are selective, depending on the cognitive demands of the task and the baseline cognitive abilities of the animals, and may be mediated, at least in part, by modulation of arousal-related neural circuits.

**Highlights:**

- FAE-20 enhanced cognitive flexibility in young adult mice
- Effects of FAE-20 were strongest in demanding cognitive tasks
- Aged poor learners benefited from FAE-20 treatment
- FAE-20 activated hypothalamic orexin neurons in male mice
- FAE-20 modulated ascending arousal systems in a sex-specific manner

## 1. Introduction

Cognitive deficits are a core feature of numerous neuropsychiatric and neurodegenerative disorders, including schizophrenia, major depressive disorder, bipolar disorder, Parkinson’s disease, and Alzheimer’s disease [1,2]. These impairments affect executive functions, cognitive flexibility, attention, working memory, and learning, and substantially contribute to reduced quality of life and poor functional outcomes. Despite the clinical relevance of these symptoms, currently available pharmacological treatments provide only limited benefits. Indeed, antidepressants, antipsychotics, and antidementia drugs often fail to produce meaningful improvements in cognition. Consequently, there remains a considerable need for novel therapeutic strategies aimed at maintaining cognitive function [3–5].

Natural products are an important resource for modern drug discovery [6,7]. In particular, medicinal plants used in traditional medicine may contain bioactive molecules with therapeutic potential that have not yet been fully characterized. Identifying and mechanistically investigating such compounds may therefore provide new opportunities for the development of cognition-enhancing treatments.

One medicinal plant that has attracted considerable scientific interest is *Rhodiola rosea* L. (Crassulaceae), commonly known as golden root. *Rhodiola rosea* has been used for centuries in traditional medicine in Asia and Eastern Europe to enhance resilience to physical and psychological stress [8,9]. Clinical and preclinical studies have reported beneficial effects of *Rhodiola rosea* extracts on fatigue, stress-related symptoms, mood, as well as learning and memory. However, the molecular constituents responsible for these effects and their mechanisms of action remain incompletely understood [10,11].

Recently, Michels and colleagues identified ferulic acid eicosyl ester (FAE-20), a naturally occurring constituent of *Rhodiola rosea* root extract, as a memory enhancing compound [12]. In their study, FAE-20 improved performance in simple Pavlovian conditioning tasks in larval and adult fruit flies (*Drosophila melanogaster)*, bees (*Apis mellifera*) and mice (*Mus musculus*). However, whether genuinely cognitive functions can be enhanced by FAE-20, as well as the neurobiological mechanisms underlying such enhancement have not yet been systematically investigated.

The present study therefore aimed to characterize the cognitive and neurobiological effects of FAE-20 in mice using a combination of behavioral and histological approaches in both young and aged animals. Cognitive flexibility was assessed using the attentional set-shifting task (ASST), whereas spatial working memory, spatial learning, and spatial memory were examined using the Y-maze and Barnes maze. In addition, neuronal activity across selected neurotransmitter systems was evaluated using c-Fos immunohistochemistry, alongside markers of hippocampal plasticity, including adult neurogenesis and dendritic spine density.

We hypothesized that FAE-20 would enhance cognitive performance, particularly in tasks requiring cognitive flexibility and spatial learning, and that these behavioral effects would be accompanied by regionand neurotransmitter system-specific alterations within circuits implicated in learning and memory, such as the cholinergic, dopaminergic, and orexinergic systems.

## 2. Methods

### 2.1. Animals

Adult male and female C57BL/6J mice were used in all experiments. Animals were obtained from the institute’s in-house breeding colony (original breeders from Charles River, Sulzfeld, Germany) and housed in same-sex groups of up to eight per cage under controlled environmental conditions (temperature: 22 ± 2 °C; humidity: 55 ± 10%). Mice were maintained on a 12 h light/dark cycle (lights on at 6:00 a.m.) and kept in standard type III cages with nesting material and shelters. Food and water were provided *ad libitum*, except during the attentional set-shifting task (ASST; see below). Behavioral testing was conducted during the light phase between 8:00 a.m. and 4:00 p.m.

All procedures were performed in accordance with the European Directive for the protection of animals used for scientific purposes (2010/63/EU) and were approved by the local authority (Landesverwaltungsamt Sachsen-Anhalt, Az. 42502-2-1618 Uni MD).

### 2.2. Attentional set-shifting task (ASST)

The ASST was used to assess cognitive flexibility in a two-choice discrimination paradigm in which mice learned to discriminate reward-predictive cues while ignoring irrelevant stimulus dimensions. The task included reversal learning and intraand ex-tra-dimensional set-shifts and was conducted according to previously published protocols [13–15].

#### 2.2.1. Apparatus and stimuli

Testing was performed in custom-built rectangular chambers (41 cm × 22 cm × 24 cm) consisting of a waiting compartment and two choice compartments separated by a transparent partition. Access was controlled by sliding doors. Each compartment contained a bowl (5 cm diameter, 2.5 cm height); the waiting compartment bowl contained water, whereas the bowls in the choice compartments were filled with digging media, one of which contained a food reward (approximately 20 mg of chocolate rice).

Stimuli were presented in two dimensions: visual/tactile (digging media) and olfactory (odorants). Six exemplars per dimension were used. Digging media varied in texture, size, and color (small and large beads, gravel, or clay granulate), while odor cues consisted of distinct odorants (valeric acid, both carvone enantiomers, eucalyptol, citral, or 2-phenylethanol). Odorants were diluted (1:20) in paraffin oil), and 30 µl were applied to filter paper attached to the bowls.

#### 2.2.2. Habituation and pre-training

Prior to testing, mice were food-restricted for one week to maintain 90–95% of their baseline body weight. Animals were habituated to the reward and apparatus over several days, including home-cage exposure to baited bowls, group habituation in the testing apparatus, and individual training sessions. During individual sessions, mice learned to retrieve rewards from bowls without discriminative cues, with the reward location varied pseudorandomly and progressively buried to promote digging behavior.

#### 2.2.3. Testing procedure and behavioral measures

Testing was conducted over four consecutive days and comprised seven phases: day 1: simple discrimination; day 2: compound discrimination and first reversal; day 3: intra-dimensional shift and second reversal; day 4: extra-dimensional shift and third reversal.

At the start of each trial, mice were placed in the waiting compartment. After opening the sliding doors, animals freely explored both choice compartments. A choice was defined as active digging, upon which the unchosen option was removed. Correct responses were rewarded, whereas incorrect choices terminated the trial without reward.

Each phase required learning of a stimulus–reward association. In simple discrimination, only one dimension was relevant. In compound discrimination, a second, irrelevant dimension was introduced. Reversal phases involved switching reward contingencies. During the intra-dimensional shift, new exemplars were introduced within the same relevant dimension, whereas in the extra-dimensional shift, the dimension of relevance changed. A phase was completed upon six consecutive correct responses. The number of trials and errors to criterion were recorded. Stimulus configurations and reward positions were pseudorandomized across animals.

### 2.3. Spontaneous alternation behavior in the Y-maze

Spontaneous alternation performance was assessed to evaluate spatial working memory, based on the natural tendency of rodents to explore novel environments. Total arm entries were additionally used as a measure of locomotor activity. The procedure followed established protocols [16,17].

#### 2.3.1. Apparatus

The Y-maze consisted of three identical arms (36 cm length, 10 cm width, 12 cm height) arranged at 120° angles, with an opaque floor and transparent walls. The maze was placed centrally in a testing room under moderate illumination (100–150 lx), with distal spatial cues available.

#### 2.3.2. Testing procedure and behavioral measures

At the beginning of the test, each mouse was placed at the distal end of one arm facing the center. Animals were allowed to freely explore the maze for 8 min in the absence of rewards. Arm entries were recorded manually and defined as entry with all four paws. After testing, mice were returned to their home cages, and the maze was cleaned with 70% ethanol between trials

Spontaneous alternations were defined as successive entries into three different arms without repetition. The alternation percentage was calculated as: (alternations) / (total arm entries − 2) × 100. Both alternation performance and total arm entries were analyzed.

### 2.4. Barnes-maze

The Barnes maze was used to assess spatial learning and memory based on the ability of mice to locate an escape box using distal cues. A previously published protocol was used [18].

#### 2.4.1. Apparatus

The maze consisted of a circular platform (92 cm diameter) with 20 evenly spaced holes (5 cm diameter). One hole led to an escape box (17 × 13 × 6 cm) located underneath the platform and containing bedding material. The position of the escape box could be changed and one of four positions were used in a balanced way for the individual mice.

Testing was conducted under bright illumination (ca. 600 lx) to create a mildly aversive environment. Distal visual cues were positioned around the maze and remained constant throughout testing. Trials were initiated from a central start position, and behavior was recorded via an overhead camera and analyzed using tracking software (EthoVision XT, Noldus). The maze and escape box were cleaned with 70% ethanol between trials.

#### 2.4.2. Experimental procedure and behavioral measures

##### 2.4.2.1. Habituation

On day 1, mice were habituated to the testing room for 1 h and subsequently placed in a transparent cylinder at the center of the maze. They were then gently guided toward the escape hole and allowed to enter the escape box, where they remained for 1 min.

##### 2.4.2.2. Training

Training was conducted over two consecutive days, with three trials per day and inter-trial intervals of at least 20 min. Each trial started with placement of the mouse in the central start cylinder. After 10 s, the cylinder was removed, and the mouse was allowed to explore the maze freely for up to 3 min. Latency to enter the escape box was recorded. If the mouse failed to locate the escape box within this time, it was gently guided to the target location. After each trial, animals remained in the escape box for 1 min.

##### 2.4.2.3. Probe test

Twenty-four hours after the final training session, a probe trial was performed in the absence of the escape box. Mice were allowed to freely explore the maze for 3 min. Spatial memory performance was assessed by measuring the latency and path length to reach the former escape location, as well as the time spent in each quadrant (zone) of the maze (target vs. mean of left, right, and opposite).

### 2.5. Immunohistochemistry

Mice were deeply anesthetized by intraperitoneal injection of ketamine hydrochloride (100 mg/kg), xylazine hydrochloride (20 mg/kg), and acepromazine maleate (3 mg/kg), and transcardially perfused with saline for 2 min followed by Zamboni’s fixative (4% paraformaldehyde, 0.2% picric acid in PBS) for 10 min. Brains were removed, post-fixed overnight, and cryoprotected in 30% sucrose. Coronal sections (40 µm) were cut using a cryostat.

Free-floating sections were processed for immunohistochemistry. Sections were rinsed in phosphate-buffered saline (PBS), incubated in 50% ethanol for 30 min, and washed three times in PBS containing 0.3% Triton X-100 (PBS-T). After a brief incubation in working buffer (PBS with 0.3% Triton X-100 and 1% normal goat serum, NGS), sections were blocked for 1 h at room temperature in blocking buffer (PBS with 0.3% Triton X-100 and 3% NGS). Sections were incubated for 72 h at room temperature with primary antibodies diluted in working buffer. Rabbit anti-c-Fos antibody (1:2000; Synaptic Systems, #226008) was co-incubated with one of the following primary antibodies: guinea pig anti-ChAT (choline acetyltransferase; 1:1000; Synaptic Systems, #297015), guinea pig anti-TH (tyrosine hydroxylase; 1:2000; Synaptic Systems, #213104), mouse anti-GAD67 (glutamic acid decarboxylase 67; 1:3000; Sigma-Aldrich, MAB5406), or mouse anti-orexin (1:300; Santa Cruz, sc-80263). A separate series of sections from aged mice was processed for doublecortin (DCX) using a rabbit anti-DBX antibody (1:2000, Abcam, Ab18723).

Following primary antibody incubation, sections were washed four times in PBS-T and incubated for 1.5 h at room temperature with a biotinylated anti-rabbit secondary antibody (1:1000 in working buffer). After three washes in PBS-T, sections were incubated for 1 h at room temperature in ABC Elite solution (Vector Laboratories; 25 µl reagent A and 25 µl reagent B per 10 ml PBS-T). Sections were then washed three times and incubated for 20 min at room temperature in biotin-tyramide solution (24 µl biotin-tyramide and 20 µl H_2_O_2_ per 10 ml PBS-T). After three additional washes, sections were incubated overnight at room temperature with Streptavidin-Cy3 (1:1000), anti-mouse-Cy5 (1:400) and anti-guinea pig Cy5 (1:400) diluted in working buffer. Finally, sections were washed in PBS, mounted onto gelatin-coated slides, air-dried, dehydrated, and coverslipped with DPX mounting medium.

For quantitative analysis, photomicrographs were obtained and analyzed using predefined rectangular regions of interest (LH: 500 × 500 µm) placed bilaterally within the center of the laterodorsal and pedunculopontine tegmental nuclei (cholinergic neurons), ventral tegmental area (dopaminergic neurons), locus coeruleus (noradrenergic neurons), lateral hypothalamus (orexinergic neurons), dorsal raphe nucleus (serotonergic neurons), and the lateral and basolateral nuclei of the amygdala (GABAergic neurons). Regions were identified with reference to a standard mouse brain atlas. Within each region, c-Fos-positive neurons, transmitter-specific neurons, and double-labeled neurons were manually counted by an experimenter blind to experimental conditions. Between two and four sections per hemisphere were analyzed. For each animal, the mean number of labeled neurons per mm² (averaged across both hemispheres) was calculated and used for statistical analysis.

### 2.6. Assessment of hippocampal dendritic spine density

Brains were fixed in 4% paraformaldehyde in 0.1 M phosphate buffer for 24 h at 4°C. Subsequently, they were transferred to 7 ml of Golgi–Cox solution (MORPHIS-TO, Offenbach, Germany) and incubated in the dark for 14 days. After 7 days, the Golgi–Cox solution was replaced with an equal volume of freshly prepared solution. The brains were then transferred to 25% sucrose in PBS and incubated at 4°C for 1–2 days. Coronal sections (100 µm) were prepared using a vibratome (Pelco Model 1000, The Vibratome Company, St. Louis, USA). Sections were mounted onto chromalaun gelatine-coated slides and allowed to dry. They were subsequently rinsed in distilled water for 2 min and incubated in 20% ammonium hydroxide prepared in distilled water (ammonium hydroxide solution, ACS reagent, 28.0–30.0% NH₃ basis; Sigma-Aldrich, Taufkirchen, Germany). Sections were then washed twice in distilled water for 2 min each. For dehydration, sections were sequentially immersed in 70%, 95%, and 100% ethanol for 5 min each. They were subsequently cleared twice in Rotihistol (Roth, Karlsruhe, Germany) for 10 min per incubation and coverslipped using DPX mounting medium (Merck, Darmstadt, Germany). After drying for 2 days, slides were stored at 4°C until analysis.

For analysis, brain sections were first imaged at 40× magnification using a monochrome SPOT digital camera (Diagnostic Instruments, USA) mounted on a Leitz DM R microscope (Leica, Wetzlar, Germany). Secondary dendrites of CA1 pyramidal neurons located in the stratum radiatum were selected for analysis. Dendritic length was measured from the primary dendrite using the NeuroJ plugin in ImageJ (version 1.4.3). Dendritic spines were then counted manually at 100× magnification. For each mouse, spine density (spines/µm) was determined from six secondary dendrites belonging to different neurons, and the mean value per animal was calculated. All analyses were performed blinded to the experimental groups.

### 2.7. Experimental procedures

The following experiments were performed

#### ASST experiment 1

Seven female and five male mice aged 2–3 months were included. Animals received intraperitoneal injections of FAE-20 at the beginning of each test day, 30 min before behavioral testing. Mice underwent the attentional set-shifting task (ASST) twice, with a one-week interval between sessions, and treatment order was pseudo-randomized.

#### ASST experiment 2

Six female and six male mice aged 2–3 months were used. Treatment begun three days before behavioral testing. Mice received daily intraperitoneal injections of either vehicle or FAE-20 during the three days preceding the ASST, as well as 30 min before testing on each ASST day. As in Experiment 1, mice underwent a second ASST session after a one-week interval and received the alternate treatment.

#### Effects of FAE-20 on brain activity and neurotransmitter systems

Female and male mice aged 2–3 months were treated with either vehicle or FAE-20 for three consecutive days (n = 6 per sex and treatment group). On the fourth day, brains were collected 1 h after the final injection and processed for immunohistochemical staining of c-Fos, a marker of neuronal activation, in combination with markers of specific neurotransmitter systems. Orexinergic neurons were identified by orexin immunolabeling, whereas tyrosine hydroxylase (TH), choline acetyltransferase (ChAT), and glutamate decarboxylase 67 (GAD67) were used as markers of catecholaminergic, cholinergic, and GABAergic neurons, respectively.

#### Y-maze experiment 1 (acute treatment)

Fourteen male mice aged 2–3 months received intraperitoneal vehicle injections, whereas thirteen mice received FAE-20. Thirty minutes later, animals were tested in the Y-maze.

#### Y-maze experiment 2 (14 days treatment)

Eight male mice aged 24–26 months received daily intraperitoneal vehicle injections for 14 days, whereas five mice received FAE-20. Thirty minutes after the final injection, mice were tested in the Y-maze. On the same day, animals were sacrified by isoflurane overdose and their brains collected for doublecortin immunohistochemistry (left hemisphere) and Golgi staining for dendritic spine density analysis (right hemisphere).

#### Barnes maze experiment 1 (young adult mice)

Eighteen mice aged 2–3 months were included. Throughout the experiment, animals received daily injections of vehicle (n = 9) or FAE-20 (n = 9) either 30 min before habituation, before the first trial on each of the two consecutive training days, on the mornings of the days between training and probe testing, and 30 min before the probe trial.

#### Barnes maze experiment 2 (aged mice)

Eleven mice aged 24 months underwent the same procedure as in Barnes maze experiment 1, with five mice receiving vehicle and six receiving FAE-20. Following a four-week washout period, mice were retested in the Barnes maze using the alternate treatment and a new escape box location. One mouse died during the washout period.

#### Barnes maze experiment 3 (aged mice, delayed application)

Fifteen mice aged 24 months underwent a modified Barnes maze procedure. During the first testing week, mice were trained without treatment. In the second week, animals were trained to locate a new escape box position and received either vehicle (n = 7) or FAE-20 (n = 8). One hour after the probe trial, mice were sacrificed and their brains processed for c-Fos and orexin immunohistochemistry.

### 2.8. Statistical Analyses

Data were analyzed using analysis of variance (ANOVA), followed by post hoc comparisons using the two-linear step-up procedure of Benjamini, Krieger and Yekutieli [19] where appropriate. Data are presented as mean ± SEM, and statistical significance was set at p < 0.05.

ASST experiment 1 and 2: repeated-measures ANOVAs were conducted with treatment and ASST phase as within-subject factors and sex as a between-subject factor. Barnes maze experiment 1, 2 and 3: for training and probe trial data, repeated-measures ANOVAs included trial or zone as within-subject factors and treatment as a between-subject factor. In Barnes maze experiment 2, treatment was analyzed as an additional within-subject factor. Y-maze experiment 1 and 2: Y-maze performance, doublecortin-positive cell counts, hippocampal spine density, and orexin/c-Fos immunohistochemistry in aged mice were analyzed using unpaired t-tests. Effects of FAE-20 on brain activity and neurotransmitter systems: two-way ANOVAs with sex and treatment as between-subject factors were used.

## 3. Results

### 3.1. Attentional set-shifting task (ASST)

#### ASST experiment 1

Figure 1 shows the mean trials to criterion following vehicle or FAE-20 treatment, averaged across all ASST phases (Fig. 1A) or displayed separately for each phase (Fig. 1B). A repeated-measures ANOVA with sex as a between-subject factor and treatment and phase as within-subject factors revealed a significant main effect of phase (F_6,60_ = 2.31, p = 0.046). Although FAE-20 treatment reduced the number of trials to criterion, the main effect of treatment did not reach statistical significance (F_1,10_ = 4.61, p = 0.058). No other main effects or interactions were significant (Fs < 1.87, ps > 0.10).

**Fig. 1.**
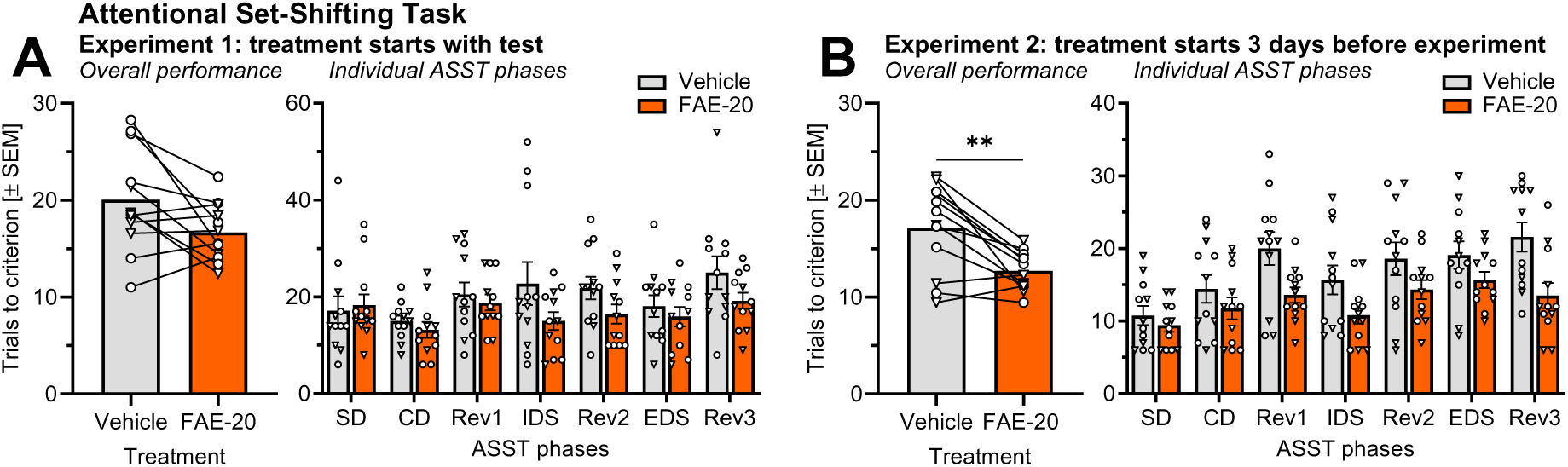
Effects of FAE-20 on ASST performance. (A) FAE-20 treatment initiated on the first day of the ASST did not significantly affect overall ASST performance. However, its effects appeared more pronounced on days 3 and 4 than on days 1 and 2. (B) When FAE-20 treatment was initiated 3 days before ASST testing, ASST performance was significantly improved. This effect was independent of ASST phase. Abbreviations: CD, compound discrimination; EDS, extra-dimensional shift; IDS, intra-dimensional shift; Rev1–3, reversals 1–3; SD, simple discrimination. **p < 0.01, main effect in ANOVA.

Based on the observation that the effects of FAE-20 appeared more pronounced during the second half of the first ASST experiment, particularly during the intra-dimensional shift, Rev2, and Rev3 phases, we hypothesized that several days of treatment might be required to produce significant behavioral outcomes. Therefore, a second ASST experiment was conducted, in which treatment began three days before behavioral testing.

Behavioral data from the second ASST experiment are shown in Fig. 1C–D. A repeated-measures ANOVA using the same factors as in the first experiment again revealed a significant main effect of phase (F_6,60_ = 7.59, p < 0.0001). Post hoc analyses showed that mice generally required more trials to criterion during the first and second reversal phases compared to the preceding compound discrimination and intra-dimensional shift phases, respectively (ts > 2.20, ps < 0.03), and that the extra-dimensional shift was more difficult than the intra-dimensional shift (t = 2.83, p = 0.006). Importantly, FAE-20 treatment significantly improved ASST performance (F_1,10_ = 15.51, p = 0.003). This effect did not differ across phases (F_6,60_ = 1.39, p = 0.24) or between sexes (F_1,10_ = 0.06, p = 0.80). There was also no main effect of sex on ASST performance (F_1,10_ = 0.004, p = 0.95).

These two ASST experiments demonstrate that FAE-20 improves cognitive flexibility, as measured by trials to criterion, and that treatment must begin several days before testing to produce significant outcomes.

### 3.2. Effects of FAE-20 on brain activity and neurotransmitter systems

The pharmacological mechanism underlying the effects of FAE-20 is currently unknown. Therefore, identifying the neurotransmitter systems affected by FAE-20 may provide an entry point to understand its mechanism of action. Towards this end, mice were treated with either vehicle or FAE-20, their brains were collected 1 h after the final injection, and subsequently processed for immunohistochemical staining of c-Fos, a marker of neuronal activation, together with markers for specific neurotransmitter systems.

A FAE-20-specific increase in c-Fos immunoreactivity would indicate activation of the respective brain region, whereas increased co-localization of c-Fos with neurotrans-mitter-specific markers would suggest activation of neurons belonging to the corresponding neurotransmitter system.

#### Lateral hypothalamus and orexin neurons (**Fig. 2A**_1_–A_6_)

The total number of orexin-positive neurons was not affected by sex or treatment, and no interaction between these factors was observed (Fig. 2A_1_; Fs < 2.74, ps > 0.11). In contrast, female mice exhibited a higher number of c-Fos-positive neurons than male mice (Fig. 2A_2_; F_1,20_ = 25.56, p < 0.0001). A similar sex difference was observed for neurons co-expressing c-Fos and orexin (Fig. 2A_3_; F_1,20_ = 32.31, p < 0.0001). No significant main effects of FAE-20 treatment were detected for either marker alone (Fs < 3.26, ps > 0.08).

**Fig 2.**
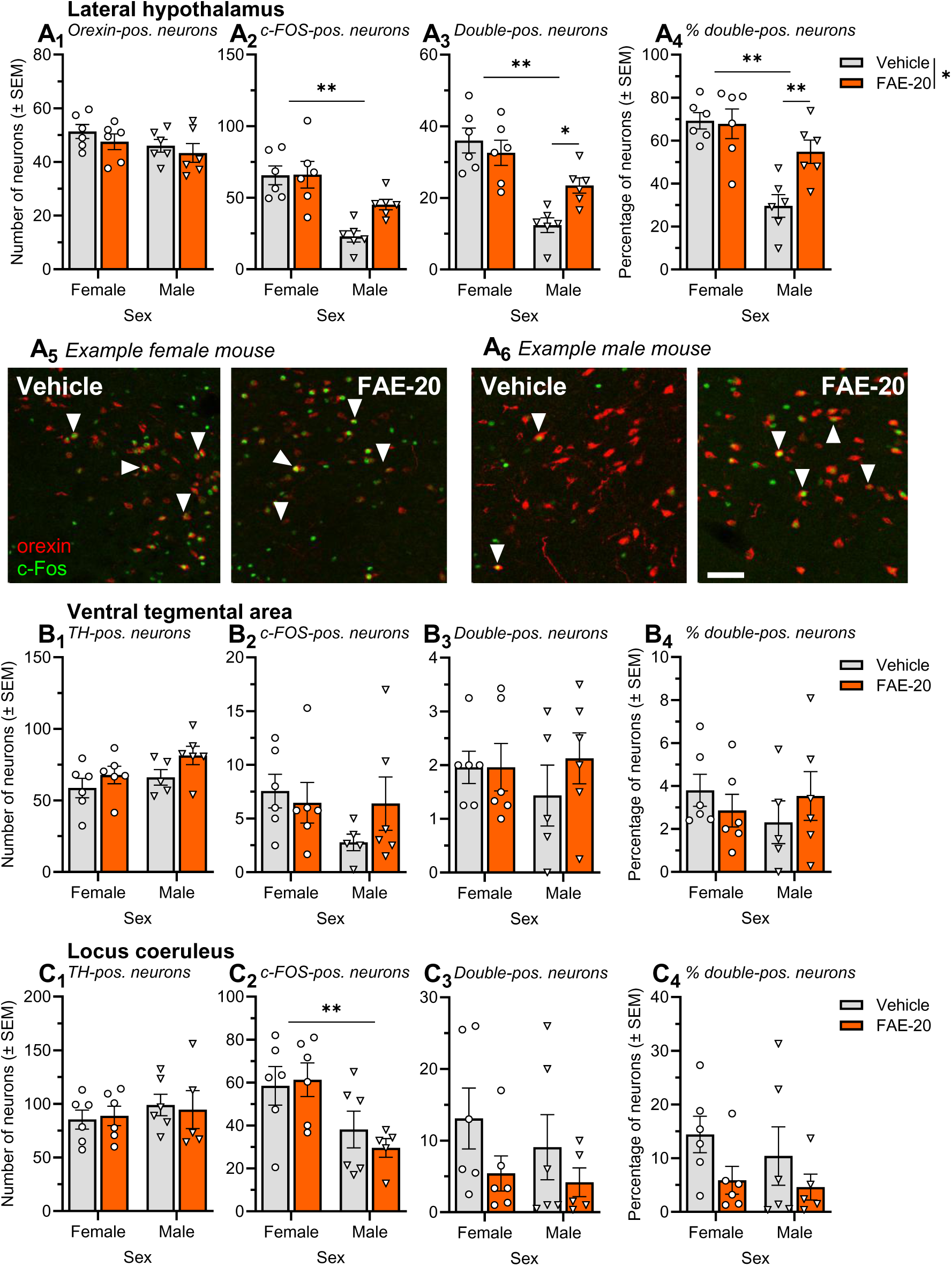
Effects of FAE-20 on brain activity and neurotransmitter systems (orexin, dopamine, noradrenaline). Depicted are the absolute or relative numbers of neurons in the (A) lateral hypothalamus, (B) ventral tegmental area, and (C) locus coeruleus. For each brain region, the first subpanel shows the number of neurons positive for the respective neurotransmitter marker, the second subpanel shows the number of c- Fos-positive neurons, and the third and fourth subpanels show the absolute and relative numbers of double-positive neurons, respectively. Panels A_5_ and A_6_ show photomicrographs from representative individual mice whose cell counts were close to the respective group means. Bar: 100 µm. Abbreviations: TH, tyrosine hydroxylase. **p < 0.01, as indicated (ANOVA followed by post hoc comparisons).

The interaction between sex and treatment approached significance for c-Fos-positive neurons (F_1,20_ = 3.02, p = 0.098) and was significant for the number of neurons co-expressing c-Fos and orexin (Fig. 2A_3_; F_1,20_ = 6.31, p = 0.02). A comparable interaction was also observed for the percentage of orexin neurons positive for c-Fos (Fig. 2A_4_-2A_6_; F_1,20_ = 6.31, p = 0.02). In addition, this analysis revealed a significant main effect of FAE-20 treatment (F_1,20_ = 4.77, p = 0.04). Post hoc comparisons showed that FAE-20 significantly increased both the absolute number and the percentage of c-Fos-positive orexin neurons in male mice (ts > 2.70, ps < 0.02), whereas no such effect was observed in female mice (ts < 0.85, ps > 0.40).

#### Ventral tegmental area and TH-positive neurons (**Fig. 2B**_1_-B_4_)

Neither the number of c-Fos-positive neurons (Fig. 2B_1_), the number of TH-positive neurons (Fig. 2B_2_), nor the absolute or relative number of neurons co-expressing c-Fos and TH (Fig. 2B_3_ and 2B_4_, respectively) in the ventral tegmental area was significantly affected by sex or FAE-20 treatment (Fs < 1.70, ps > 0.20). Likewise, no significant interactions between sex and treatment were observed (Fs < 1.60, ps > 0.22).

#### Locus coeruleus and TH-positive neurons (**Fig. 2C**_1_-C_4_)

In the locus coeruleus, the only significant effect observed was a main effect of sex on the number of c-Fos-positive neurons (Fig. 2C_2_; F1,19 = 10.72, p = 0.004), with female mice showing higher numbers than male mice. Notably, there were trend-level main effects of FAE-20 treatment on both the absolute and relative numbers of neurons co-expressing c-Fos and TH (Fig. 2C_3_; F1,19 = 3.03, p = 0.098 and Fig. 2C_4_; F1,19 = 3.59, p = 0.07, respectively), indicating reduced numbers following FAE-20 treatment. No other main effects or interactions were significant (Fs < 0.74, ps > 0.40).

#### Laterodorsal tegmental nucleus and ChAT-positive neurons (**Fig. 3A**_1_-3A_6_)

Significant main effects of sex and FAE-20 treatment, as well as a significant interaction between both factors, were observed for the number of ChAT-positive neurons in the laterodorsal tegmental nucleus (Fig. 3A_1_; Fs > 13.02, ps < 0.002). Post hoc analysis revealed a significant FAE-20-induced increase in female mice (t = 7.87, p < 0.0001).

**Fig 3.**
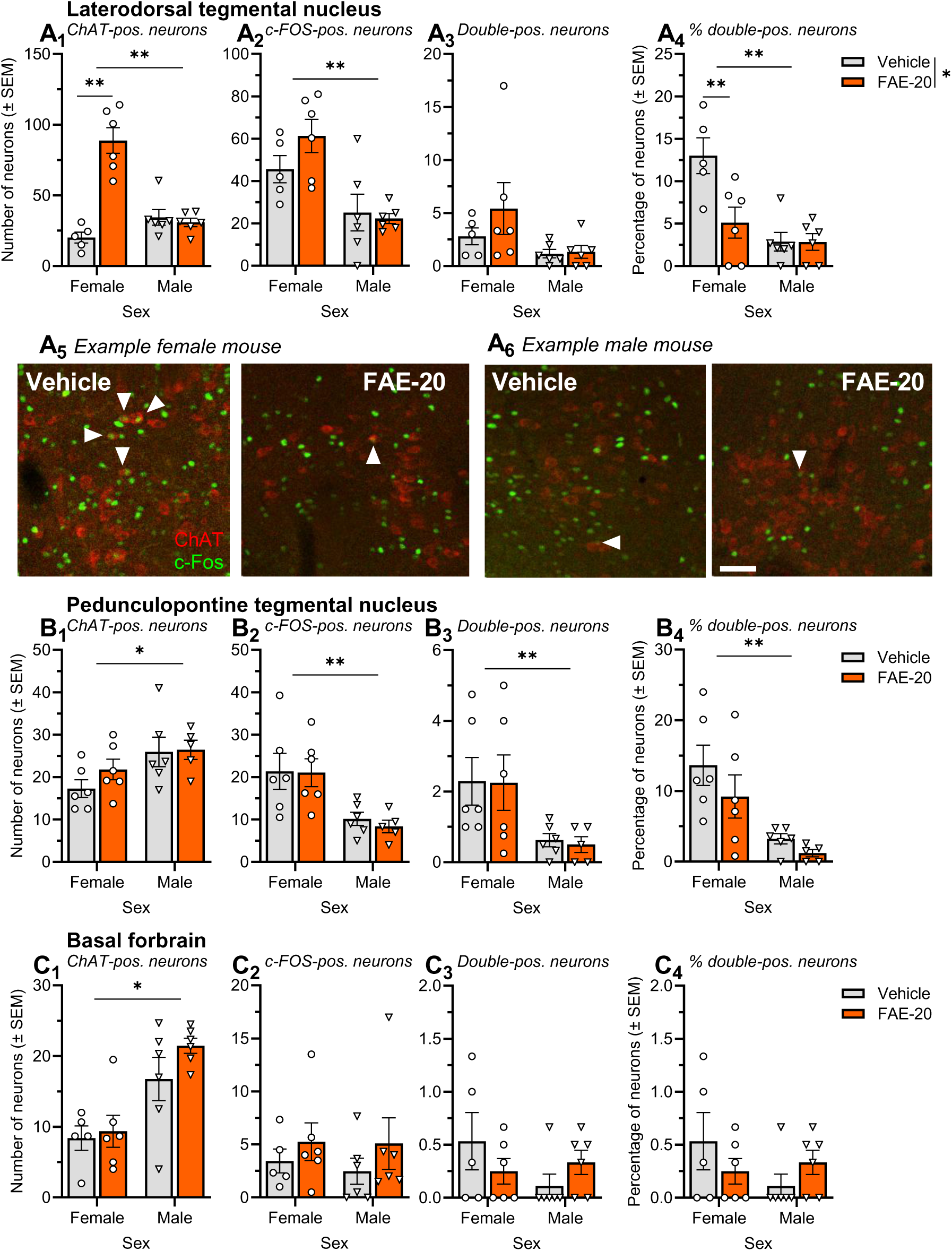
Effects of FAE-20 on brain activity and neurotransmitter systems (acetyl choline). Depicted are the absolute or relative numbers of neurons in the (A) laterodorsal tegmental nucleus, (B) ventral tegmental area, and (C) locus coeruleus. For each brain region, the first subpanel shows the number of neurons positive for the respective neurotransmitter marker, the second subpanel shows the number of c-Fos-positive neurons, and the third and fourth subpanels show the absolute and relative numbers of double-positive neurons, respectively. Panels A_5_ and A_6_ show photomicrographs from representative individual mice whose cell counts were close to the respective group means. Bar: 100 µm. Abbreviations: ChAT, choline acetyltransferase. *p < 0.05, **p < 0.01, as indicated (ANOVA followed by post hoc comparisons).

For c-Fos-positive neurons (Fig. 3A_2_), there was a significant main effect of sex, with higher numbers in female mice (F_1,19_ = 18.78, p = 0.0004), but no significant effect of treatment or interaction (Fs < 1.83, ps > 0.19).

For the absolute number of ChAT/c-Fos double-positive neurons (Fig. 3A_3_), the interaction between sex and treatment did not reach significance (F_1,19_ = 4.25, p = 0.053). However, for the percentage of ChAT neurons co-expressing c-Fos (Fig. 3A_4_-3A_6_), this interaction was significant (F_1,19_ = 6.60, p = 0.02), accompanied by significant main effects of sex and treatment (Fs > 6.70, ps < 0.02). Post hoc comparisons showed a significant reduction in the percentage of double-positive neurons in female mice following FAE-20 treatment (t = 3.56, p = 0.002).

#### Pedunculopontine tegmental nucleus and ChAT-positive neurons (**Fig. 3B**_1_-3B_4_)

In the pedunculopontine tegmental nucleus, only significant main effects of sex were observed. The number of ChAT-positive neurons was increased in male mice (Fig. 3B_1_; F_1,19_ = 6.22, p = 0.02), whereas the number of c-Fos-positive neurons, as well as the absolute and relative numbers of ChAT/c-Fos double-positive neurons, were decreased in male mice (Fig. 3B_2_–3B_4_; Fs > 6.21, ps < 0.03). No effects of FAE-20 treatment and no sex × treatment interactions were detected (Fs < 2.07, ps > 0.16).

#### Basal forebrain and ChAT-positive neurons (**Fig. 3C**_1_-3C_4_)

As in the previous brain region, the basal forebrain showed a higher number of ChAT-positive neurons in male mice (Fig. 3C_1_; F_1,19_ = 21.33, p = 0.0002). No other main effects or interactions were observed in any of the remaining analyses (Fs < 2.62, ps > 0.12).

#### Basolateral amygdala and GAD67-positive neurons (**Fig. 4A**_1_-4A_4_)

For GAD67-positive neurons, significant main effects were observed for both sex and FAE-20 treatment (Fig. 4A_1_; F_1,20_ = 5.87, p = 0.03 and F_1,20_ = 4.47, p = 0.047, respectively), whereas the interaction between these factors was not significant (F_1,20_ = 0.46, p = 0.50). No significant effects were detected in the analysis of c-Fos-positive neurons (Fig. 4A_2_). The absolute and relative numbers of GAD67/c-Fos double-positive neurons were reduced in male mice (Fig. 4A_3_; F_1,20_ = 2.50, p = 0.005 and Fig. 4A_4_; F_1,20_ = 5.27, p = 0.03), while no effects of sex or interactions were observed (Fs < 2.50, ps > 0.12).

**Fig 4.**
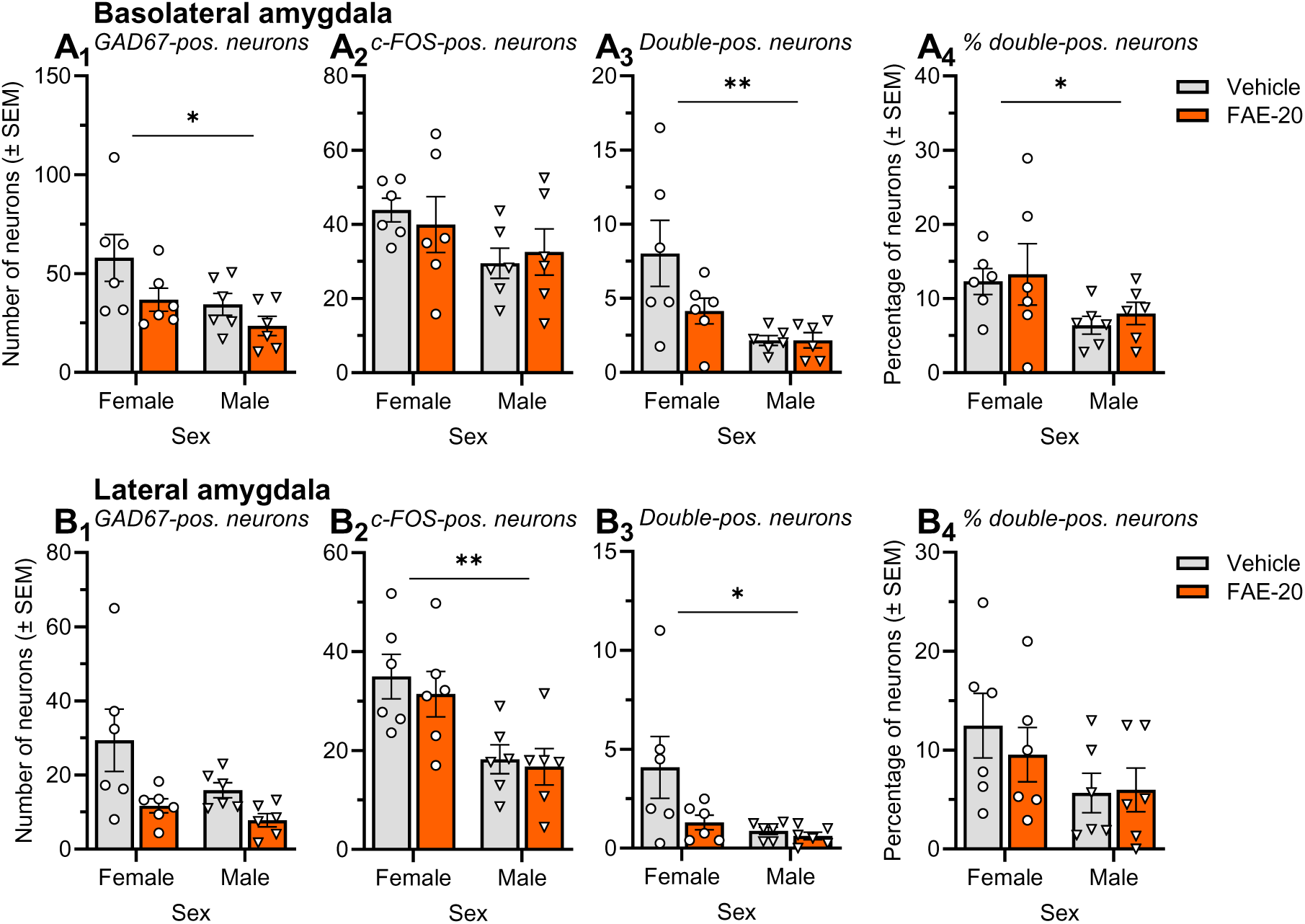
Effects of FAE-20 on brain activity and neurotransmitter systems (GABA). Depicted are the absolute or relative numbers of neurons in the (A) basolateral amygdala, and (B) lateral amygdala. For each brain region, the first subpanel shows the number of neurons positive for the respective neurotransmitter marker, the second subpanel shows the number of c-Fos-positive neurons, and the third and fourth subpanels show the absolute and relative numbers of double-positive neurons, respectively. Abbreviations: GAD67, 67-kDa glutamate decarboxylase. *p < 0.05, **p < 0.01, as indicated (ANOVA followed by post hoc comparisons).

**Fig. 5.**
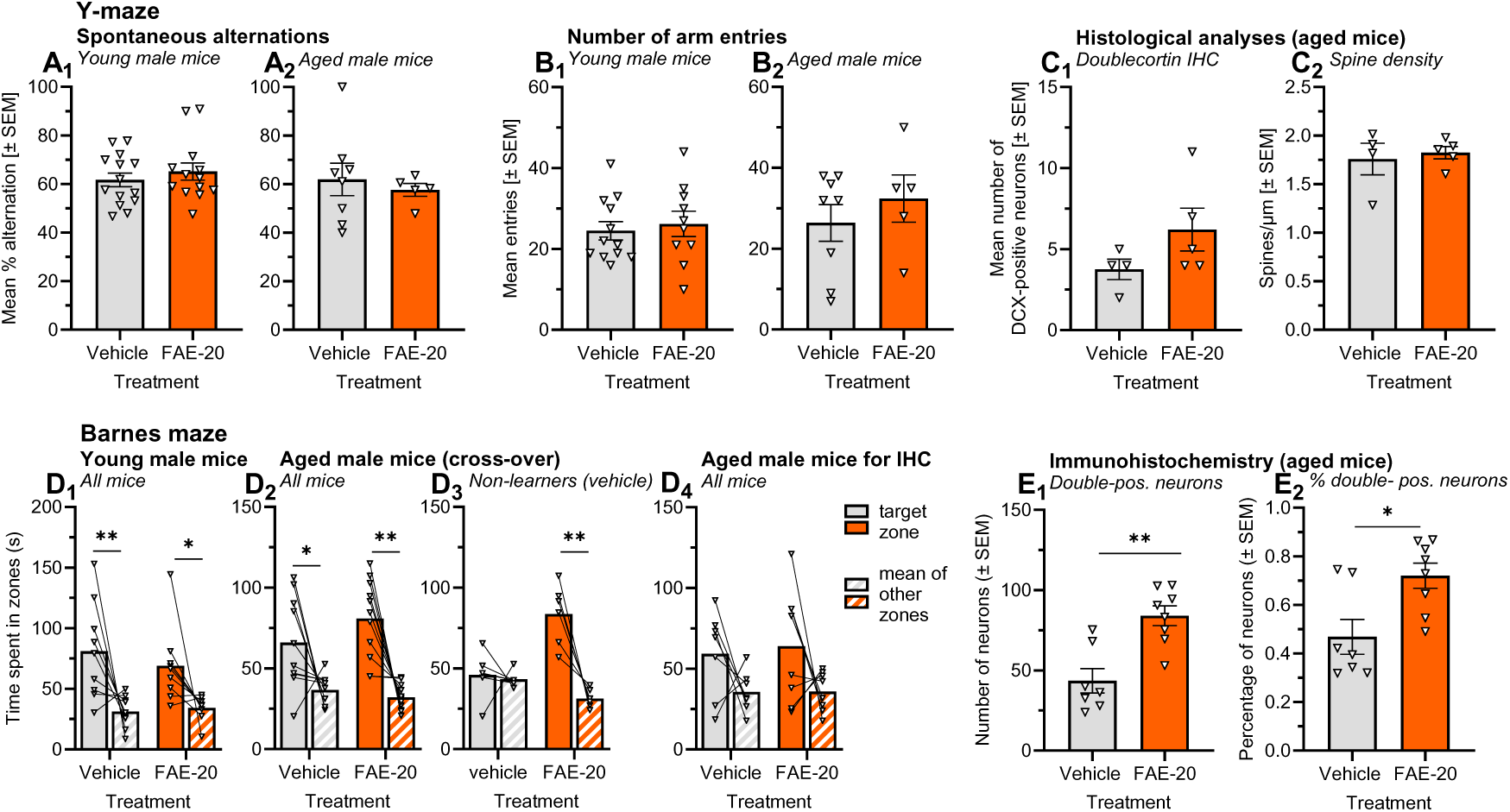
Effects of FAE-20 on behavior in the Y-maze and Barnes maze. FAE-20 treatment did not affect the percentage of spontaneous alternations (A) or the number of arm entries (B) in young (A_1_, B_1_) or aged (A_2_, B_2_) mice. In aged mice, FAE-20 also had no effect on the number of doublecortin-positive neurons (C_1_) or hippocampal spine density (C_2_). Similarly, no treatment effects were observed in the Barnes maze in young (D_1_) or aged (D_2_, D_4_) mice. However, among mice exhibiting absent or poor spatial learning under vehicle treatment, FAE-20 significantly improved spatial learning performance (D_3_). Furthermore, FAE-20 increased both the absolute and relative numbers of c-Fos/orexin double-positive neurons in aged male mice (E_1_–E_2_). Abbreviations: IHC, immunohistochemistry. *p < 0.05, **p < 0.01, as indicated (ANOVA followed by post hoc comparisons).

#### Lateral amygdala and GAD67-positive neurons (**Fig. 4B**_1_-4B_4_)

The number of GAD67-positive neurons was reduced following FAE-20 treatment (Fig. 4B_1_; F_1,20_ = 8.15, p = 0.01), with a trend toward lower numbers in male mice (F_1,20_ = 3.71, p = 0.07). This male-associated reduction was significant for the number of c-Fos-positive neurons (Fig. 4B_2_; F_1,20_ = 15.57, p = 0.0008) and for the absolute number of GAD67/c-Fos double-positive neurons (Fig. 4B_3_; F_1,20_ = 5.74, p = 0.03), while only a trend was observed for the relative number of double-positive neurons (Fig. 4B_4_; F_1,20_ = 3.97, p = 0.06). In addition, a trend-level effect of FAE-20 treatment was detected for the absolute number of double-positive neurons (Fig. 4B_3_; F_1,20_ = 3.97, p = 0.06). No other main effects or interactions were significant (Fs < 2.40, ps > 0.13).

Overall, FAE-20 induced sex-, regionand cell type–specific alterations in neuronal activity across the neurotransmitter systems examined. In male mice, FAE-20 increased the activity of orexinergic neurons in the lateral hypothalamus, whereas in female mice it reduced the activity of cholinergic neurons in the laterodorsal tegmental nucleus. Measures of dopaminergic neurons in the ventral tegmental area and noradrenergic neurons in the locus coeruleus were largely unaffected. Independent of treatment, sex differences were observed in the number of cholinergic and GABAergic neurons, as well as in the activity of tegmental and amygdaloid neurons.

### 3.3. Effects of FAE-20 on behavior in the Y-maze and Barnes maze

The Y-maze and Barnes maze are well-established paradigms for assessing relatively simple forms of spatial working memory and spatial learning. Because young mice typically perform well in these tasks, enhancing treatment effects can be difficult to detect. Therefore, both young and aged mice were included. As only male aged mice were available, these experiments were conducted exclusively in males. Following behavioral testing, additional histological analyses were performed in the aged mice.

#### Behavior in the Y-maze

Spontaneous alternation behavior, a measure of spatial working memory, was not affected by FAE-20 treatment in either young (Fig. 3A_1_; t = 0.77, p = 0.45) or aged mice (Fig. 3A_2_t = 0.63, p = 0.63). Similarly, the number of arm entries remained unchanged in both young (Fig. 3B_1_; t = 0.45, p = 0.66) and aged animals (Fig. 3B_2_; t = 0.82, p = 0.43), indicating that FAE-20 did not exert nonspecific effects on locomotor activity.

#### Doublecortin immunohistochemistry

In aged mice, FAE-20 treatment did not alter the number of doublecortin-positive neurons (Fig. 3C_1_; t = 1.54, p = 0.17), suggesting that FAE-20 had no effect on adult hippocampal neurogenesis.

#### Spine densitity in the dorsal hippocampus

The density of dendritic spines in the dorsal hippocampus was not affected by 14 days of FAE-20 treatment (Fig. 3C_2_; t = 0.42, p = 0.69).

#### Behavior in the Barnes maze

Both young and aged mice reduced the distance to the escape box over the course of training, indicating successful spatial learning (young: F_5,80_ = 92, p = 0.0001; aged: F_5,50_ = 3.07, p = 0.02; data not shown). No main effects of treatment or treatment × trial interaction were observed in either age group (Fs < 1.61, ps > 0.16).

Successful spatial learning was confirmed in the probe trial, as mice spent significantly more time in the target zone, where the escape box had been located during training, than in the other zones (young: Fig. 3D_1_, F_1,16_ = 14.17, p = 0.002; aged: Fig. 3D_2_, F_1,38_ = 40.70, p > 0.0001). Again, no main effects of FAE-20 treatment or treatment × zone interaction were detected (Fs < 2.51, ps > 0.12).

Notably, the experiment in aged mice was conducted using a crossover design, in which all animals received both treatments, separated by a washout period. This design allowed a separate analysis of mice that failed to show spatial learning under vehicle treatment (Fig. 3D_3_). In this poor-learner subgroup, significant main effects of zone (F_1,10_ = 18.35, p = 0.002) and a zone × treatment interaction (F_1,10_ = 15.00, p = 0.003) were observed, suggesting that FAE-20 may improve spatial learning and memory in those aged mice that show poor baseline performance under control conditions.

#### Orexin immunohistochemistry

A separate cohort of aged male mice received subchronic FAE-20 treatment and underwent Barnes maze training followed by a probe trial. These animals were subsequently, i.e. without cross-over treatment, used for c-Fos and orexin immunohistochemistry 1 h after the probe trial. Similar to the previous cohort, Barnes maze learning was observed (zones: F_1,13_ = 5.25, p = 0.04), whereas FAE-20 treatment had no effect on performance (treatment × zones: F_1,13_ = 0.04, p = 0.85; data not shown).

Importantly, c-Fos and orexin immunohistochemistry in aged mice replicated the effects of FAE-20 previously observed in young animals. FAE-20 treatment increased both the absolute and the relative number of neurons co-expressing c-Fos and orexin (Fig. 3E_1_, t = 4.19, p = 0.001; Fig. 3E_2_, t = 2.91, p = 0.01), indicating enhanced activation of orexinergic neurons in the lateral hypothalamus.

Taken together, FAE-20 did not affect spatial working memory in the Y-maze, spatial learning and memory in the Barnes maze, locomotor activity, hippocampal neurogenesis, or dendritic spine density in young or aged male mice. However, a subgroup analysis revealed improved Barnes maze performance in aged mice with poor baseline learning under vehicle treatment. In addition, FAE-20 consistently increased the activation of orexinergic neurons in the lateral hypothalamus of aged mice, replicating findings previously observed in young animals (Fig. 2).

## 4. Discussion

The present study aimed to characterize the cognitive and neurobiological effects of the *Rhodiola rosea* constituent FAE-20 using a combination of behavioral and histological approaches in young and aged mice. The main findings can be summarized as follows. First, FAE-20 improved performance in the ASST, indicating enhanced cognitive flexibility, provided that treatment was initiated several days before testing. Second, FAE-20 did not affect spatial working memory-based spontaneous alternation behavior in the Y-maze or overall spatial learning in the Barnes maze in either young or aged mice. However, a subgroup analysis in aged animals revealed improved Barnes maze performance in those mice that failed to acquire the task under vehicle treatment. Third, histological analyses identified sex-, region and cell type-specific effects of FAE-20 on neuronal activity within components of the ascending arousal system. Specifically, FAE-20 increased the activity of orexinergic neurons in the lateral hypothalamus of male mice, whereas it reduced the activity of cholinergic neurons in the LDTg of female mice. In contrast, no consistent FAE-20 effects were observed on hippocampal neurogenesis, dendritic spine density, dopaminergic neurons of the ventral tegmental area, noradrenergic neurons of the locus coeruleus, or GABAergic neurons in the amygdala.

The behavioral findings suggest that the cognition-enhancing effects of FAE-20 depend strongly on the cognitive demands of the task and on the baseline cognitive abilities of the tested animals. The most robust effects were observed in the ASST, a paradigm that requires animals to acquire, reverse, and shift attention across multiple stimulus dimensions [20,21]. Compared with many standard rodent learning paradigms, the ASST places substantially higher demands on executive functions and cognitive flexibility and is therefore particularly sensitive to pharmacological manipulations affecting higher-order cognitive processes. Consistent with this interpretation, our laboratory has repeatedly detected cognitive effects of pharmacological interventions in the ASST in young animals, both under normal conditions [13] and following the induction of mild cognitive deficits [14,22]. The significant improvement observed after subchronic FAE-20 treatment therefore suggests that the compound facilitates executive aspects of cognition rather than producing nonspecific changes in activity or motivation.

In contrast, FAE-20 did not affect performance in the Y-maze. This finding is not necessarily inconsistent with the ASST results. Spontaneous alternation behavior primarily reflects short-term spatial working memory and is generally considered a relatively simple task for healthy young mice [18,23]. Importantly, the use of working memory in this paradigm is entirely spontaneous, as animals do not experience any disadvantage for ’failing’ to alternate between arms. Consequently, the task places only limited cognitive demands on the animals. Even in aged mice, performance often remains sufficiently high to reduce the sensitivity of the paradigm for detecting cognition-enhancing effects. Similar considerations apply to the Barnes maze [24]. Both young and aged animals successfully acquired the task and showed robust preference for the target zone during the probe trial. Under such conditions, ceiling effects may substantially reduce the likelihood of detecting treatment-related improvements.

The use of aged animals in the Y-maze and Barnes maze experiments was motivated by the expectation that enhancements would be easier to detect in animals with age-related low performance. Unfortunately, only male aged mice were available for these experiments. Consequently, the findings from these paradigms cannot be generalized to female animals, representing a limitation of the present study.

Particularly noteworthy is the Barnes maze crossover experiment in aged mice. The crossover design allowed each animal to serve as its own control and enabled the identification of a subgroup of animals that exhibited little or no learning under vehicle treatment. Within this subgroup, FAE-20 significantly improved spatial learning and memory performance. This finding is of interest because cognition-enhancing compounds are often expected to exert their largest effects under conditions of impaired cognitive function rather than in healthy subjects performing near ceiling levels [25,26]. The present results therefore suggest that FAE-20 may be especially effective in individuals with reduced baseline cognitive performance, in agreement with our previous findings [12]. Moreover, the crossover approach employed here may represent a valuable experimental strategy for future studies investigating FAE-20 and other putative cognitive enhancers. Detecting improvements in subjects with low baseline performance may provide a more sensitive approach than relying solely on group-level analyses in cognitively intact populations.

The histological findings provide first clues regarding the neurobiological mechanisms underlying the behavioral effects of FAE-20 in mice. Although the molecular target of the compound remains unknown, the observed alterations in neuronal activity suggest that FAE-20 influences specific components of the ascending arousal system, a network critically involved in the regulation of wakefulness, attention, motivation, and cognitive performance.

In male mice, FAE-20 increased the proportion of activated orexinergic neurons in the lateral hypothalamus. Importantly, a similar effect was observed in both young and aged animals, supporting the robustness of this finding. Orexin neurons play a central role in promoting wakefulness and behavioral arousal and project extensively to cortical and subcortical regions involved in cognitive control [27–29]. Increased orexinergic activity is associated with enhanced attention, motivation, vigilance, and prefrontal cortical activation, all of which can facilitate learning and executive functioning [14,17,30–33]. The observed activation of orexin neurons therefore provides a plausible mechanistic explanation for the beneficial effects of FAE-20 on cognitive flexibility and potentially on learning in cognitively impaired aged animals.

The effects observed in female mice were less intuitive but may nevertheless be relevant for cognitive performance. FAE-20 reduced the proportion of activated cholinergic neurons within the laterodorsal tegmental nucleus. At first glance, this finding appears inconsistent with the established literature linking enhanced cholinergic neurotransmission in the cortex and hippocampus to improved cognition [34,35]. However, cholinergic neurons in the brainstem and midbrain fulfill functions that differ substantially from those of basal forebrain cholinergic projections. In particular, activation of cholinergic neurons within the laterodorsal tegmental nucleus has frequently been associated with stress-related responses and increased behavioral impulsivity [36–39]. Excessive activation of this system may therefore interfere with optimal cognitive performance [40–42]. Conversely, reduced activity within these neurons may decrease impulsive responding and facilitate executive functions such as cognitive flexibility. Thus, despite their opposite direction, the effects observed in female mice may converge functionally with the orexinergic activation observed in males by ultimately promoting cognitive performance, though through distinct neural pathways.

Notably, FAE-20 did not alter markers of adult hippocampal neurogenesis or dendritic spine density in aged mice. These findings suggest that the behavioral effects of FAE-20 are unlikely to depend on major structural alterations of hippocampal plasticity, at least within the treatment duration investigated here. Instead, the available evidence points toward modulation of neuronal activity within arousal- and attention-related networks as a more likely mechanism. Similarly, only minor or inconsistent effects were observed within dopaminergic, noradrenergic, and GABAergic systems, indicating a certain degree of neurochemical specificity.

Several limitations of the present study should be acknowledged. Sample sizes were relatively small in the aged cohorts. Furthermore, the mechanistic conclusions are based on correlational c-Fos analyses and therefore do not establish causal relationships between the observed neuronal activation patterns and the behavioral effects of FAE-20. Future studies should combine pharmacological, chemogenetic, or optogenetic approaches with behavioral testing to directly determine whether orexinergic or cholinergic signaling mediates the observed beneficial effects of FAE-20. In addition, the molecular target of FAE-20 remains unknown and will require further investigation.

In conclusion, the present findings provide evidence that FAE-20 possesses cognition-enhancing properties. Beneficial effects were observed primarily in cognitively demanding tasks and in animals with poor baseline cognitive performance, suggesting that the compound may be particularly effective under conditions of cognitive impairment. Moreover, the study identifies two potential, sex-specific neurobiological mechanisms involving orexinergic and cholinergic components, respectively, of the ascending arousal system. Although these findings do not reveal the target of FAE-20, they provide an important starting point for future mechanistic investigations. Overall, FAE-20 emerges as a promising candidate for the development of novel therapeutic approaches aimed at improving cognitive function when it is compromised, warranting further preclinical and translational studies.

## Acknowledgements

The authors would like to thank Dr. Christian König-Bethke for preparing the test solutions, Kathrin Freke for her support with animal care, and Uwe Diesterheft for his assistance in developing and refining the behavioral setups.

## CRediT authorship contribution statement

**Dana Mayer:** Formal analysis, Investigation, Visualization. **Iman Hassan:** Formal analysis, Investigation. **Firat Taskaya:** Formal analysis, Investigation. **Afsana Chowdhury:** Investigation. **Evelyn Kahl:** Formal analysis, Investigation. **Thomas Endres:** Conceptualization, Supervision, Writing – review & editing. **Volkmar Leßmann:** Resources, Writing – review & editing. **Bertram Gerber:** Conceptualization, Resources, Writing – review & editing**. Markus Fendt:** Conceptualization, Data curation, Formal analysis, Funding acquisition, Resources, Supervision, Visualization, Writing – original draft, Writing – review & editing.

## Data statement

Data will be made available on request.

## 5.#References

[1] R.A. McCutcheon, R.S.E. Keefe, P.K. McGuire, Cognitive impairment in schizophrenia: aetiology, pathophysiology, and treatment, Mol Psychiatry 28 (2023) 1902–1918. 10.1038/s41380-023-01949-9.

[2] J. Hugo, M. Ganguli, Dementia and cognitive impairment: epidemiology, diagnosis, and treatment, Clin Geriatr Med 30 (2014) 421–442. 10.1016/j.cger.2014.04.001.

[3] M.J. Millan, Y. Agid, M. Brüne, E.T. Bullmore, C.S. Carter, N.S. Clayton, R. Connor, S. Davis, B. Deakin, R.J. DeRubeis, B. Dubois, M.A. Geyer, G.M. Goodwin, P. Gorwood, T.M. Jay, M. Joëls, I.M. Mansuy, A. Meyer-Lindenberg, D. Murphy, E. Rolls, B. Saletu, M. Spedding, J. Sweeney, M. Whittington, L.J. Young, Cognitive dysfunction in psychiatric disorders: characteristics, causes and the quest for improved therapy, Nat Rev Drug Discov 11 (2012) 141–168. 10.1038/nrd3628.

[4] A. Vita, W. Gaebel, A. Mucci, G. Sachs, S. Barlati, G.M. Giordano, G. Nibbio, M. Nordentoft, T. Wykes, S. Galderisi, European Psychiatric Association guidance on treatment of cognitive impairment in schizophrenia, Eur Psychiatry 65 (2022) e57. 10.1192/j.eurpsy.2022.2315.

[5] Z. Arvanitakis, R.C. Shah, D.A. Bennett, Diagnosis and management of dementia: review, Jama 322 (2019) 1589–1599. 10.1001/jama.2019.4782.

[6] A.G. Atanasov, S.B. Zotchev, V.M. Dirsch, C.T. Supuran, Natural products in drug discovery: advances and opportunities, Nat Rev Drug Discov 20 (2021) 200–216. 10.1038/s41573-020-00114-z.

[7] D.J. Newman, G.M. Cragg, Natural products as sources of new drugs over the nearly four decades from 01/1981 to 09/2019, J Nat Prod 83 (2020) 770–803. 10.1021/acs.jnatprod.9b01285.

[8] E. Ivanova Stojcheva, J.C. Quintela, The efectiveness of Rhodiola rosea L. preparations in alleviating various aspects of life-stress symptoms and stress-induced conditions-encouraging clinical evidence, Molecules 27 (2022). 10.3390/molecules27123902.

[9] Y. Li, V. Pham, M. Bui, L. Song, C. Wu, A. Walia, E. Uchio, F. Smith-Liu, X. Zi, Rhodiola rosea L.: an herb with anti-stress, anti-aging, and immunostimulating properties for cancer chemoprevention, Curr Pharmacol Rep 3 (2017) 384–395. 10.1007/s40495-017-0106-1.

[10] A. Panossian, Understanding adaptogenic activity: specificity of the pharmacological action of adaptogens and other phytochemicals, Ann N Y Acad Sci 1401 (2017) 49–64. 10.1111/nyas.13399.

[11] A. Coors, M. Brosch, E. Kahl, R. Khalil, B. Michels, A. Laub, K. Franke, B. Gerber, M. Fendt, Rhodiola rosea root extract has antipsychotic-like effects in rodent models of sensorimotor gating, J. Ethnopharmacol. 235 (2019) 320–328. 10.1016/j.jep.2019.02.031.

[12] B. Michels, H. Zwaka, R. Bartels, O. Lushchak, K. Franke, T. Endres, M. Fendt, I. Song, M. Bakr, T. Budragchaa, B. Westermann, D. Mishra, C. Eschbach, S. Schreyer, A. Lingnau, C. Vahl, M. Hilker, R. Menzel, T. Kähne, V. Leßmann, A. Dityatev, L. Wessjohann, B. Gerber, Memory enhancement by ferulic acid ester across species, Sci. Adv 4 (2018) eaat6994.

[13] S. Afzal, N. Dürrast, I. Hassan, E. Soleimanpour, P.L. Tsai, D.C. Dieterich, M. Fendt, Probing cognitive flexibility in Shank2-deficient mice: Effects of D-cycloserine and NMDAR signaling hub dynamics, Prog. Neuropsychopharmacol. Biol. Psychiatry 134 (2024) 111051. 10.1016/j.pnpbp.2024.111051.

[14] J. Duske, N. D’Souza, D. Mayer, D.C. Dieterich, M. Fendt, Orexinergic modulation of chronic jet lag-induced deficits in mouse cognitive flexibility, Neuropsychopharmacology 50 (2025) 762–771. 10.1038/s41386-024-02017-8.

[15] L. Seifried, E. Soleimanpour, D.C. Dieterich, M. Fendt, Cognitive flexibility in mice: effects of puberty and role of NMDA receptor subunits, Cells 12 (2023). 10.3390/cells12091212.

[16] M. Dembeck, D.C. Dieterich, M. Fendt, The GluN2C/D-specific positive allosteric modulator CIQ rescues delay-induced working memory deficits in mice, Behav. Brain Res. 456 (2024) 114716. 10.1016/j.bbr.2023.114716.

[17] A. Burckhardt, A. Rakowsky, E. Kahl, S. Morchhale, D. Mayer, N. Panagiotou, L. Permien, N. Faesel, M. Fendt, Differential effects of orexin system activation on dizocilpine-induced schizophrenia-like behaviors in mice, Neuropeptides 116 (2026) 102587. 10.1016/j.npep.2026.102587.

[18] E.A.K. Prieur, N.M. Jadavji, Assessing spatial working memory using the spontaneous alternation Y-maze test in aged male mice, Bio Protoc 9 (2019) e3162. 10.21769/BioProtoc.3162.

[19] Y. Benjamini, A.M. Krieger, D. Yekutieli, Adaptive linear step-up procedures that control the false discovery rate, Biometrika 93 (2006) 491–507. 10.1093/biomet/93.3.491.

[20] D.S. Tait, E.A. Chase, V.J. Brown, Attentional set-shifting in rodents: a review of behavioural methods and pharmacological results, Curr. Pharm. Des. 20 (2014) 5046–5059. 10.2174/1381612819666131216115802

[21] G.B. Bissonette, E.M. Powell, Reversal learning and attentional set-shifting in mice, Neuropharmacology 62 (2012) 1168–1174. 10.1016/j.neuropharm.2011.03.011

[22] M.A.B. Siddik, M. Fendt, D-cycloserine rescues scopolamine-induced deficits in cognitive flexibility in rats measured by the attentional set-shifting task, Behav. Brain Res. 431 (2022) 113961. 10.1016/j.bbr.2022.113961.

[23] A.K. Kraeuter, P.C. Guest, Z. Sarnyai, The Y-maze for assessment of spatial working and reference memory in mice, Methods Mol Biol 1916 (2019) 105–111. 10.1007/978-1-4939-8994-2_10.

[24] K. Gawel, E. Gibula, M. Marszalek-Grabska, J. Filarowska, J.H. Kotlinska, Assessment of spatial learning and memory in the Barnes maze task in rodents-methodological consideration, Naunyn Schmiedebergs Arch Pharmacol 392 (2019) 1–18. 10.1007/s00210-018-1589-y.

[25] R. de Jongh, I. Bolt, M. Schermer, B. Olivier, Botox for the brain: enhancement of cognition, mood and pro-social behavior and blunting of unwanted memories, Neurosci. Biobehav. Rev. 32 (2008) 760–776. 10.1016/j.neubiorev.2007.12.001.

[26] S. Granon, F. Passetti, K.L. Thomas, J.W. Dalley, B.J. Everitt, T.W. Robbins, Enhanced and impaired attentional performance after infusion of D1 dopaminergic receptor agents into rat prefrontal cortex, J Neurosci 20 (2000) 1208–1215. 10.1523/jneurosci.20-03-01208.2000.

[27] C. Alexandre, M.L. Andermann, T.E. Scammell, Control of arousal by the orexin neurons, Curr. Opin. Neurobiol. 23 (2013) 752–759. 10.1016/j.conb.2013.04.008.

[28] E. Arrigoni, T. Mochizuki, T.E. Scammell, Activation of the basal forebrain by the orexin/hypocretin neurones, Acta Physiol (Oxf) 198 (2010) 223–235. 10.1111/j.1748-1716.2009.02036.x.

[29] N. Nevárez, L. de Lecea, Recent advances in understanding the roles of hypocretin/orexin in arousal, affect, and motivation, F1000Res 7 (2018). 10.12688/f1000research.15097.1.

[30] C.B. Calva, H. Fayyaz, J.R. Fadel, Effects of intranasal orexin-A (hypocretin-1) administration on neuronal activation, neurochemistry, and attention in aged rats, Front Aging Neurosci 11 (2019) 362. 10.3389/fnagi.2019.00362.

[31] M. Stanojlovic, J.P. Pallais Yllescas, Jr., V. Mavanji, C. Kotz, Chemogenetic activation of orexin/hypocretin neurons ameliorates aging-induced changes in behavior and energy expenditure, Am J Physiol Regul Integr Comp Physiol 316 (2019) R571–r583. 10.1152/ajpregu.00383.2018.

[32] M. Stanojlovic, J.P. Pallais, M.K. Lee, C.M. Kotz, Pharmacological and chemogenetic orexin/hypocretin intervention ameliorates Hipp-dependent memory impairment in the A53T mice model of Parkinson’s disease, Mol Brain 12 (2019) 87. 10.1186/s13041-019-0514-8.

[33] A. Durairaja, S. Pandey, E. Kahl, M. Fendt, Nasal administration of orexin A partially rescues dizocilpine-induced cognitive impairments in female C57BL/6 J mice, Behav. Brain Res. 450 (2023) 114491. 10.1016/j.bbr.2023.114491.

[34] M.E. Hasselmo, The role of acetylcholine in learning and memory, Curr. Opin. Neurobiol. 16 (2006) 710–715. 10.1016/j.conb.2006.09.002.

[35] J. Haam, J.L. Yakel, Cholinergic modulation of the hippocampal region and memory function, J Neurochem 142 Suppl 2 (2017) 111–121. 10.1111/jnc.14052.

[36] S.S. Fernandes, A.P. Koth, G.M. Parfitt, M.F. Cordeiro, C.S. Peixoto, A. Soubhia, F.P. Moreira, C.D. Wiener, J.P. Oses, E. Kaszubowski, D.M. Barros, Enhanced cholinergic-tone during the stress induce a depressive-like state in mice, Behav. Brain Res. 347 (2018) 17–25. 10.1016/j.bbr.2018.02.044.

[37] C.R. Romero-Leguizamón, K.A. Kohlmeier, Stress-related endogenous neuropeptides induce neuronal excitation in the laterodorsal tegmentum, Eur. Neuropsychopharmacol. 38 (2020) 86–97. 10.1016/j.euroneuro.2020.07.008.

[38] G.L. Forster, C.D. Blaha, Laterodorsal tegmental stimulation elicits dopamine efflux in the rat nucleus accumbens by activation of acetylcholine and glutamate receptors in the ventral tegmental area, Eur. J. Neurosci. 12 (2000) 3596–3604. 10.1046/j.1460-9568.2000.00250.x.

[39] F.S. Polli, K.A. Kohlmeier, Prenatal nicotine alters development of the laterodorsal tegmentum: Possible role for attention-deficit/hyperactivity disorder and drug dependence, World J. Psychiatry 12 (2022) 212–235. 10.5498/wjp.v12.i2.212.

[40] M. Shabani, M. Ilaghi, R. Naderi, M. Razavinasab, The hyperexcitability of laterodorsal tegmentum cholinergic neurons accompanies adverse behavioral and cognitive outcomes of prenatal stress, Sci. Rep. 13 (2023) 6011. 10.1038/s41598-023-33016-2.

[41] H. Janickova, O. Kljakic, K. Rosborough, S. Raulic, S. Matovic, R. Gros, L.M. Saksida, T.J. Bussey, W. Inoue, V.F. Prado, M.A.M. Prado, Selective decrease of cholinergic signaling from pedunculopontine and laterodorsal tegmental nuclei has little impact on cognition but markedly increases susceptibility to stress, Faseb J. 33 (2019) 7018–7036. 10.1096/fj.201802108R.

[42] C.P. Lisgaras, H.E. Scharfman, The cholinergic system exerts opposing effects on memory at different stages of disease progression in Alzheimer’s and Down syndrome model systems, Alzheimers Dement. 22 (2026) e71154. 10.1002/alz.71154.

